# Data processing of product ion spectra: Methods to control false discovery rate in compound search results for non-targeted metabolomics

**DOI:** 10.1101/2024.06.16.599235

**Authors:** Fumio Matsuda

## Abstract

In non-targeted metabolomics utilizing high-resolution mass spectrometry, several database search methods have been used to comprehensively annotate the acquired product ion spectra. Recent advancements in various *in silico* prediction techniques have facilitated compound searches by scoring the degree of coincidence between a query product ion spectrum and a compound in a compound database. Certain search results may be false positives, thus necessitating a method for controlling the false discovery rate (FDR). This study proposed two simple methods for controlling the FDR in compound search results. In the pseudo-target decoy method, the FDR can be estimated without creating a separate decoy database by treating such as the positive ion mode spectra as targets and converting the negative ion mode spectra as decoys. Further, the second-rank method uses the score distribution of the second-ranked hits from the compound search as an approximation of the false-positive distribution of the top-ranked hits. The performance of these methods was evaluated by annotating the product ion spectra from

MassBank using the SIRIUS 5 CSI:Finger ID scoring method. The results indicated that the second-rank method was closer to the true FDR of 0.05. When applied to the four human metabolomics datasets, the second-rank method provided more conservative FDR estimations than the pseudo-target-decoy method. These methods enabled the identification of metabolites not present in human metabolome databases. Overall, this study demonstrates the utility of these simple methods for FDR control in non-targeted metabolomics, facilitating more reliable compound identification and the potential discovery of novel metabolites.

## Introduction

In non-targeted metabolomics utilizing high-resolution mass spectrometry, researchers employ several database search methods to comprehensively annotate the acquired product ion spectra^1)^. Molecular formula searches are performed using accurate *m*/*z* values of the precursor ions^2)^. The spectral similarity search approach has been employed to estimate the compound structure using measured spectra databases such as MassBank (Table 1)^3)^. Recent advancements in various *in silico* analysis techniques have facilitated the scoring of the degree of coincidence between a query product ion spectrum and a compound in a compound database. Several software packages that implement these methods, including MS-FINDER^4)^, SIRIUS^5,6)^, MetFrag^7,8)^, and CFM-ID^9,10)^, have been developed in recent years (Table 1). These software packages generate a ranked list of compound scores for each queried product ion spectrum. In this study, the top-ranked compound in the list was referred to as a “hit.” A compound search task is performed for every product ion spectrum in the metabolomic dataset. Consequently, because certain hits may be false positives, a score threshold must be set to control the false discovery rate (FDR) within an acceptable range^11)^.

**Table 1.**
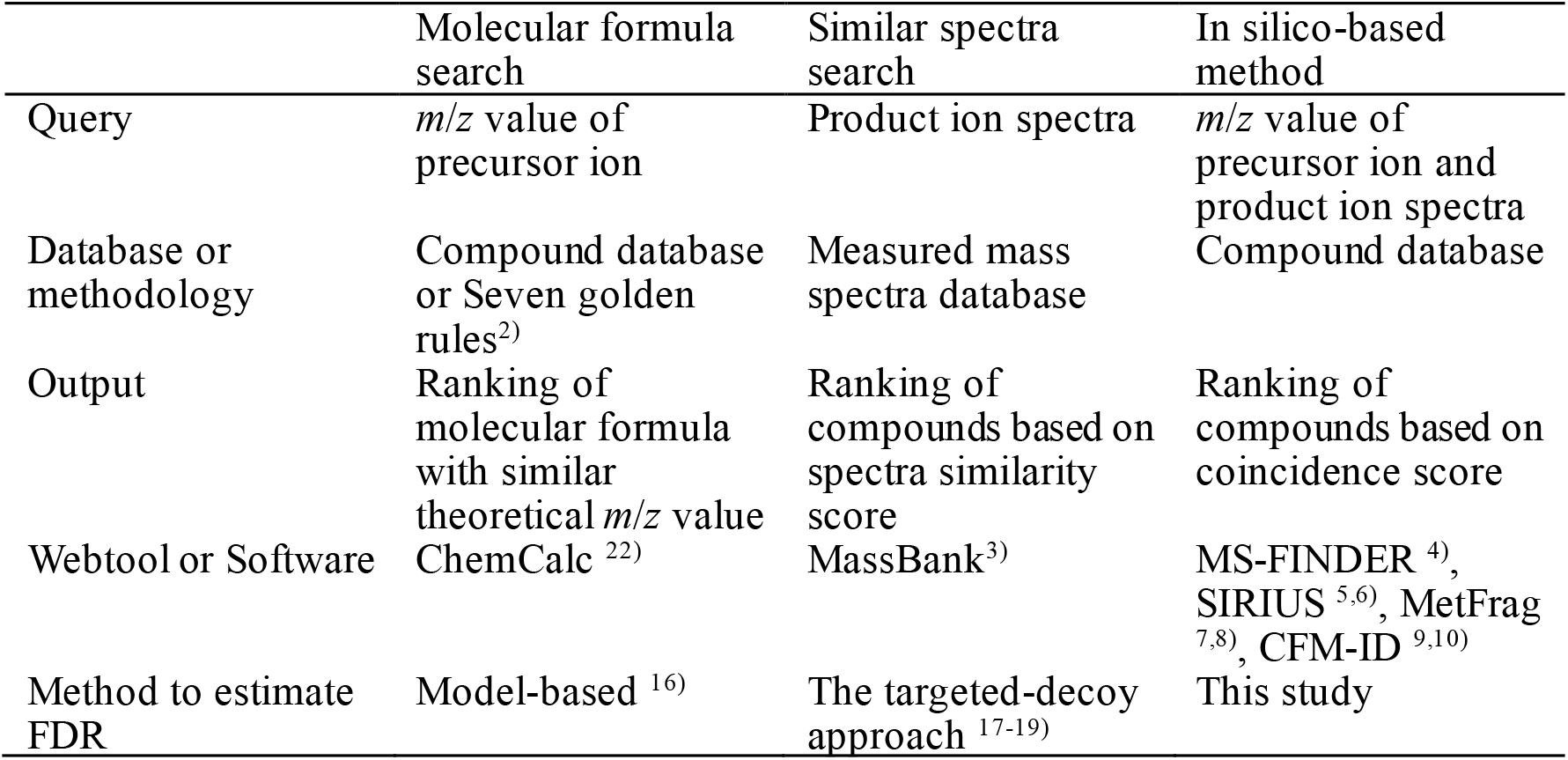
Database searching methods for structural annotation of product ion spectra

|  | Molecular formula search | Similar spectra search | In silico-based method |
| --- | --- | --- | --- |
| Query | $m/z$ value of precursor ion | Product ion spectra | $m/z$ value of precursor ion and product ion spectra |
| Database or methodology | Compound database or Seven golden rules<sup>2)</sup> | Measured mass spectra database | Compound database |
| Output | Ranking of molecular formula with similar theoretical $m/z$ value | Ranking of compounds based on spectra similarity score | Ranking of compounds based on coincidence score |
| Webtool or Software | ChemCalc<sup>22)</sup> | MassBank<sup>3)</sup> | MS-FINDER<sup>4)</sup>, SIRIUS<sup>5,6)</sup>, MetFrag<sup>7,8)</sup>, CFM-ID<sup>9,10)</sup> |
| Method to estimate FDR | Model-based<sup>16)</sup> | The targeted-decoy approach<sup>17-19)</sup> | This study |

In proteomics, the FDR of peptide identification results is estimated using the target decoy method (Fig. 1a)^12,13)^. Product ion spectra are comprehensively acquired from trypsin-digested peptides using data-dependent acquisition (DDA) mode of liquid chromatography-tandem mass spectrometry (LC/MS/MS). Peptide identification is accomplished by searching each product ion spectrum against target and decoy peptide databases ^14)^. The target database comprises all trypsin-digested peptides derived from all genome-encoded protein sequences, whereas the decoy database includes reversed sequences. The FDR is estimated as FDRs = *D*/*T* or FDRc = 2*D*/(*D*+*T*), where *T* and *D* represent the number of hits against the target and decoy databases, respectively, at a particular score-threshold level^15)^.

**Figure 1.**
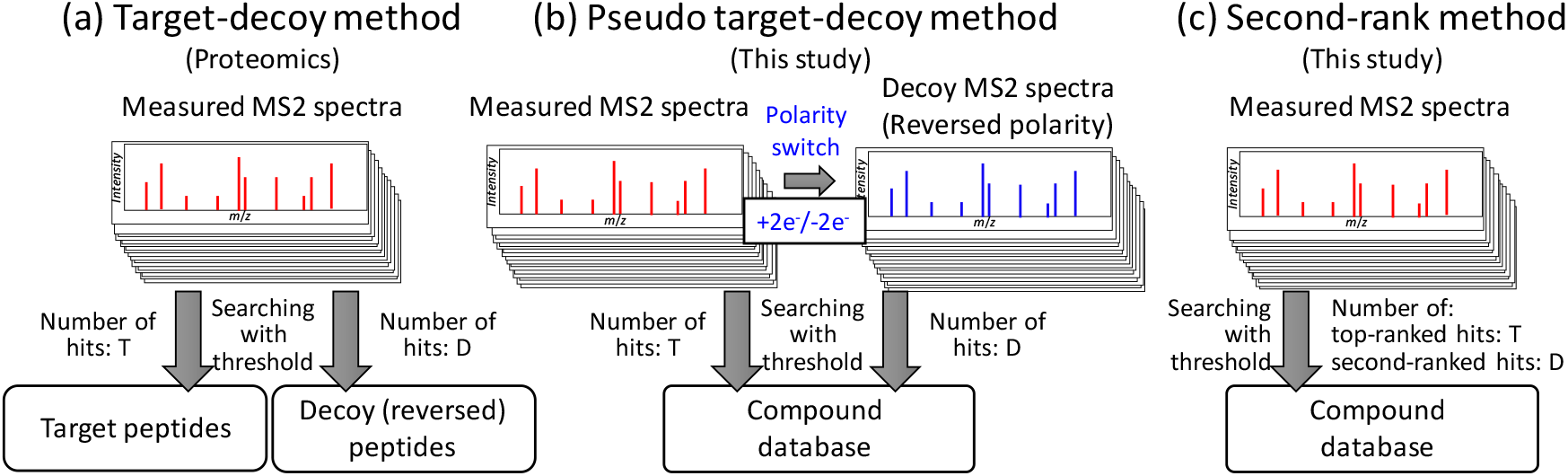
Methods to estimate false discovery rate (FDR) in product ion spectra-based peptide and compound identification results. (a) Target-decoy method used for peptide identification. (b) Pseudo target-decoy method for compound identification developed in this study. (c) Second-rank method for compound identification developed in this study.

Because we still do not know the complete list of human metabolites for compound annotation in metabolomics, there exists no established method for creating a reasonable decoy database with the same characteristics as the target database. As an alternative to the complete target decoy method, methods employing theoretical models have been reported for molecular formula searches^16)^. For spectral similarity searches, several target-decoy approaches have been reported using fragmentation tree-based^17)^, ion entropy and accurate entropy-based ^18)^, and violating the octet rule of chemistry-based^19)^ methods to create decoy spectra databases (Table 1). A Gaussian mixture model-based framework for estimating the false discovery rate (FDR) was also proposed for gas chromatography-electron impact-MS data^20)^. Recently, SIRIUS 5 implemented a new scoring method called the confidence score, which proposes that hits with a score of 0.65 or higher correspond to an FDR of 0.1 ^21)^. In addition to these methods, various other methodologies should be investigated to estimate FDR in compound search results.

In this study, the pseudo-target-decoy method and second-rank method were proposed as methods to estimate the FDR in compound search results. First, the performances of the two methods were validated using the measured product ion spectra from MassBank. We then applied the two methods to human metabolome datasets and investigated metabolites not included in the human metabolome database (HMDB).

## Experimental procedures

### Software for compound search and data processing

SIRIUS 5 (version 5.8.5) ^5,6)^ was downloaded from https://bioinformatik.uni-jena.de/software/sirius/. Compound search tasks were performed using the settings shown in Data S1. MSFINDER (version 3.61)^4)^ was downloaded from https://github.com/systemsomicslab/MsdialWorkbench/releases. Compound search tasks were performed using the settings shown in Data S1. MetFrag CL (version 2.4.5)^7,8)^ was downloaded from https://ipb-halle.github.io/MetFrag. Compound search tasks were performed using the settings shown in Data S1. Three compound databases including the yeast metabolome database (YMDB, 16042 compounds, https://www.ymdb.ca/)^23)^, human metabolome database (HMDB 220,945 compounds,https://www.hmdb.ca^24)^, and Biodatabase were used. The Biodatabase is an edited database of biological compounds available in SIRIUS 5. Although the detailed number is unclear, the number of compounds is considerably larger than that in HMDB because the Biodatabase includes biological compounds derived from HMBD, PubChem, and other compound databases. All data pre- and postprocessing tasks were performed using an in-house Python script.

### Performance test using MassBank records

All MassBank records were downloaded from the MassBank Consortium GitHub page (https://github.com/MassBank/, downloaded on 15/4/2023). First, 10,711 spectra were selected by the following criteria: AC$MASS_SPECTROMETRY: MS_TYPE = MS2, AC$INSTRUMENT_TYPE includes TOF or FT, AC$MASS_SPECTROMETRY: IONIZATION = ESI, AC$MASS_SPECTROMETRY: FRAGMENTATION_MODE = CID or no description, MS$FOCUSED_ION: PRECURSOR_TYPE = [M+H]+ or [M-H]-, CH$COMPOUND_CLASS does not include “Environmental Standard,” “Surfactant,” “Non-natural,” and “Non-Natural,” and MS$FOCUSED_ION: PRECURSOR_M/Z < 850, and the number of product ions >= 3. Second, the selected records were divided into four datasets. From the 574 compounds, several spectra were commonly acquired in both the positive and negative ion modes and designated as the CommonPos (3,388 spectra) and CommonNeg (3,100 spectra) datasets, respectively. The OthersPos (2,648 spectra) and OthersNeg (1,575 spectra) datasets comprised the remaining positive and negative ion mode data, respectively.

For each query product ion spectrum, the three software packages used in this study output a ranked list of compound scores. In this study, the top-ranked compound in the list was referred to as a hit. Even in case of multiple top-ranked compounds with the same score, they were considered hits. When one of the top-ranked compounds was correct, the hit was considered a true positive hit. For the pseudo-target-decoy method, decoy product ion spectra were generated by the addition or deduction of mass corresponding to two elections to the product ion spectra obtained in the positive and negative ion modes, respectively. To calculate the FDR, the number of hits above a given threshold level was determined for the search results from the target (*T*) and decoy (*D*) datasets. Two methods were used to calculate FDRs: FDRs = *D*/*T* and FDRc = 2*D*/(*D*+*T*)^15)^. In the second-rank method, the second-largest score in the ranked list of compound scores was used as the second-ranked score. The FDR was estimated using FDR2nd = *D*/*T*, where *D* and *T* represent the number of hits above a given threshold level for the 1^st^ and 2^nd^ ranked scores.

### Compound search and determination of false discovery rate

Four human metabolome DDA datasets were downloaded from the Metabolomics Workbench ^25)^ and MetaboLights repositories^26,27)^. Averaged spectra were generated from the product ion spectra of each dataset using the spectral averaging method described in our previous study. ^28)^ For each averaged product ion spectrum, the averaged spectra of the corresponding MS1 spectra were generated using the same method. In addition to the protonated or deprotonated molecules, additional ion forms were considered in the compound search if the corresponding ions were observed in the averaged MS1 spectra.

## Results

### Pseudo target-decoy method

To facilitate the application of the target-decoy approach without creating a decoy compound database, we focused on the fact that metabolomics employs both positive and negative ion modes for data acquisition. This indicates that a single-compound database can be used for compound searching in both ion modes. For instance, the original product ion spectra acquired in the positive ion mode (targets) can be searched against a compound database to produce target hits. The original product ion spectra could then be regarded as decoy product ion spectra acquired in the negative ion mode by adding the mass of two electrons (2e^-^) to all *m*/*z* values. Decoy hits were obtained by searching these decoy product ion spectra against an identical compound database in negative ion mode (Fig. 1b).

The proposed method was not an ideal target-decoy approach. The decoy product ion spectra do not have identical properties, because the decoy data must be generated from a compound with a different molecular formula from the target spectra. Furthermore, the fragmentation mechanisms differ between the positive and negative ion modes. Hence, we termed this method as the pseudo-target-decoy approach.

To verify the performance of this method, we utilized a dataset of the measured product ion spectra stored in the MassBank database. From MassBank records, this study used product ion spectra (MS2) obtained from the [M+H]^+^ and [M-H]^-^ of natural products by collision-induced dissociation (CID) and high-resolution mass analyzers (detailed procedure presented in Materials and Methods). The selected MassBank records were further divided into four datasets. The CommonPos (3,388 spectra) and CommonNeg (3,100 spectra) datasets included spectra commonly acquired from identical 574 compounds both in the positive and negative ion modes. The remaining positive and negative ion mode data were designated as the OthersPos (2,648 spectra) and OthersNeg (1,575 spectra) datasets, respectively.

First, the properties of the CommonPos and CommonNeg datasets were compared because these spectra were obtained from the same compounds. A comparison of the frequency distributions of *m*/*z* values, intensities, and number of product ions revealed that the CommonPos and CommonNeg datasets had similar properties, except for a slightly smaller number of product ions in the negative ion mode (Fig. S1). Next, compound search tasks were performed for each of the 3,388 spectra in the CommonPos dataset (Data S2). The compound search using SIRIUS 5 with the CSI:FingerID scoring method successfully provided a hit for 2,439 query spectra (the compound database was HMDB, and other search conditions used default values; Data S1). Each hit was checked against the correct answer and classified as a true or false positive hit. The true and false positive hits exhibited distinct score distributions (Fig. 2a), indicating that the score threshold to control FDR of 0.05 was −10.27, with 907 hits falling within this range. In this study, FDRmes represents the true FDR measured by true and false positive hits. Similar distributions were obtained for the CommonNeg dataset (Fig. S1). Moreover, similar distributions were observedeven when using a smaller database, YMDB, or a largerdatabase, Biodatabase (Fig. S1). These results indicate that therewas a positive-negativesymmetry in the compound search results using SIRIUS 5 with the CSI:FingerID scoring method. However, distinct distributions between true and false hits, or positive-negative symmetry, could not be confirmed using other software packages (Fig. S2). Therefore, subsequent analyses used SIRIUS 5 with the CSI:FingerID scoring method.

**Figure 2.**
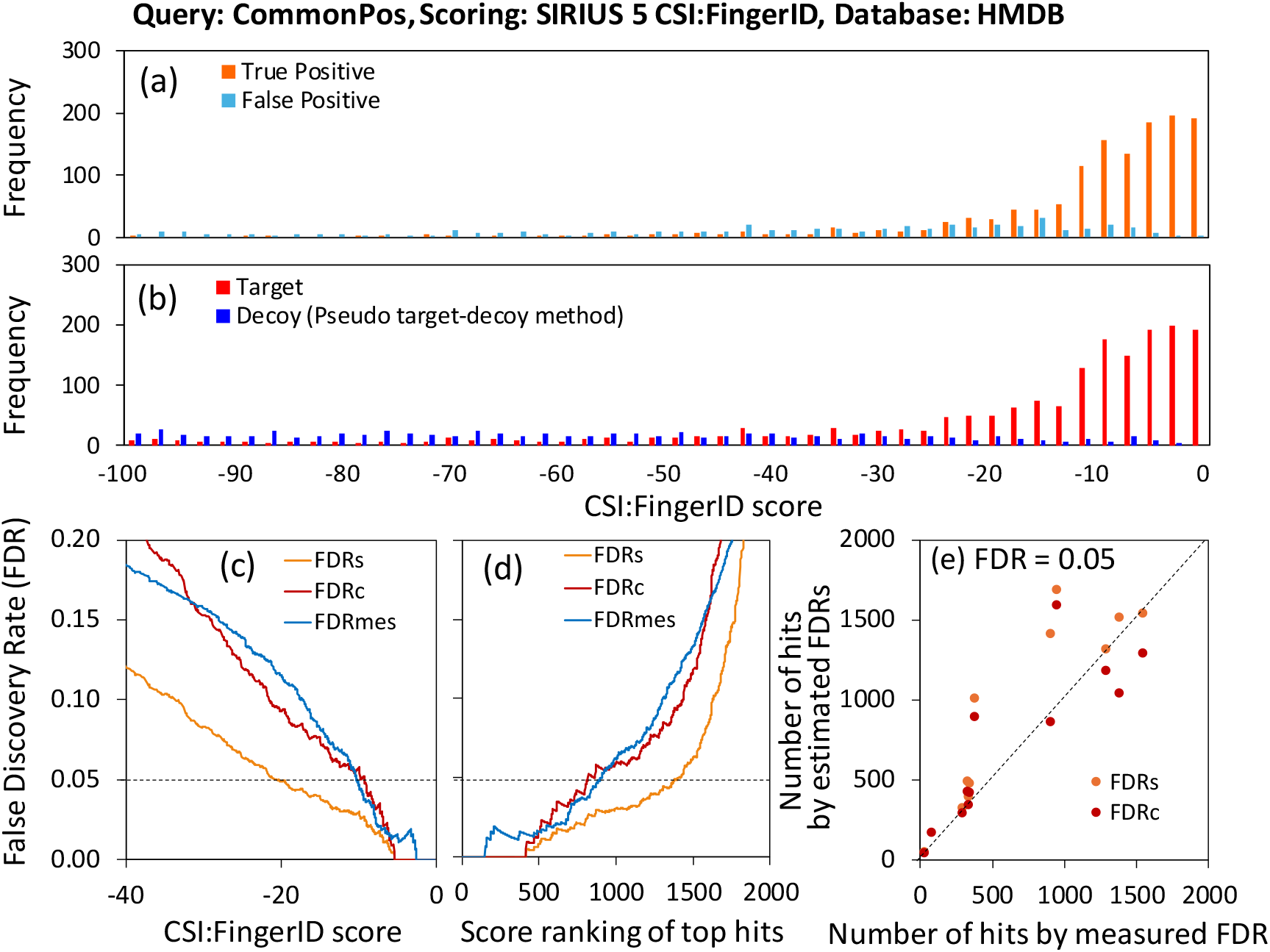
Performance evaluation of the pseudo target-decoy method. (a) Score distribution of true and false positive hits in the compound identification result. Total 3388 high-resolution mass spectra data obtained at positive ion mode were collected from MassBank. The CommonPos dataset were served for the compound search by the SIRIUS 5 CSI:Finger ID scoring method using HMDB as the compound database. (b) Score distribution of searching results of target and decoy CommonPos datasets. (c) Relationship between the CSI:Finger ID score and FDR levels. FDRmes indicate true FDR levels measured from the number of true and false hits in panel a. FDRs and FDRs represent estimated FDR levels from the number of target (*T*) and decoy (D) hits in panel b. FDRs = *D*/*T*, FDRc = 2*D*/(*T*+*D*) (d) Relationship between the score ranking of top hits and FDR levels. (e) Comparison between the number of hits determined by the true FDR (FDRmes) and estimated FDR (FDRs and FDRc).

To test the pseudo-target-decoy method, the mass of two electrons was added to all *m*/*z* values of the 3,388 spectral data points in the CommonPos dataset to produce a decoy spectral dataset. The compound search in the negative ion mode using SIRIUS 5 with the CSI:FingerID scoring method produced 2,105 hits (Data S2), with the frequency distribution shown in Figure 2b. Using the score distributions, two FDR indices, FDRs = *D*/*T* and FDRc 2*D*/(*T*+*D*), were calculated and compared with FDRmes (Fig. 2c, d); where *T* and *D* indicate the number of hits for the target and decoy datasets, respectively. The comparison showed certain similarities between FDRmes and FDRc (Fig. 2c, d). For example, the true number of hit compounds for FDRmes = 0.05 was 1,387. In contrast, the numbers estimated by FDRs and FDRc were 1,510 and 1,040, respectively, providing reasonable estimations (Fig. 2d). A similar trend was observed for the CommonNeg dataset (Fig. S3). Using the four datasets (CommonPos, CommonNeg, OhtersPos, and OthersNeg) and three compound databases of different sizes (YMDB, HMDB, BioDB), the number of hit compounds were compared among all 12 combinations at FDR = 0.05. The results showed that FDRc often underestimated the FDR in certain cases (Fig. 2e, Table S1). In addition, similar overestimation and underestimation were observed for the SIRIUS 5 with confidence score method (Table S1).

### Second-rank method

Next, we examined another approach. As mentioned previously, the top-ranked hits included both true and false positives. Each compound search also provided a second-ranked hit in addition to the top-ranked hit. Certain of the second-ranked hits had high scores that were very close to the top-ranked hits and could be considered failed attempts of false positive hits. Thus, the distribution of high scores in the second-ranked hits should be similar to that of false positives in the top-ranked hits (Fig. 1c). In peptide identification using proteomics, the score distribution of second-ranked hits is considered similar to that of false positive^14)^.

To test the second-rank method, the scores of second-ranked hits were obtained using the SIRIUS 5 CSI:Finger ID scoring method for the CommonPos dataset (Fig. 3a, Data S3). Treating these scores as decoys, score thresholds and numbers of hits were determined by FDRrank2 = *D*/*T* and compared with those from FDRmes (Fig. 3b, c). The results showed that the estimation by FDRrank2 deviated significantly from that by FDRmes, particularly in the low-score region. However, FDRrank2 provided a better approximation at FDR=0.05. For instance, the number of hits estimated by FDRrank2 was 1,205, which is a good approximation of the 1,387 hits obtained from FDRmes. In contrast, FDRrank3, calculated using third-ranked data, significantly underestimated the FDR. A comparison across all 12 combinations of the four datasets and three compound databases at FDR=0.05 (Fig. 3d) revealed that the estimation by FDRrank2 using the second-rank method was better than that using the pseudo-target-decoy method (Fig. 2e).

**Figure 3.**
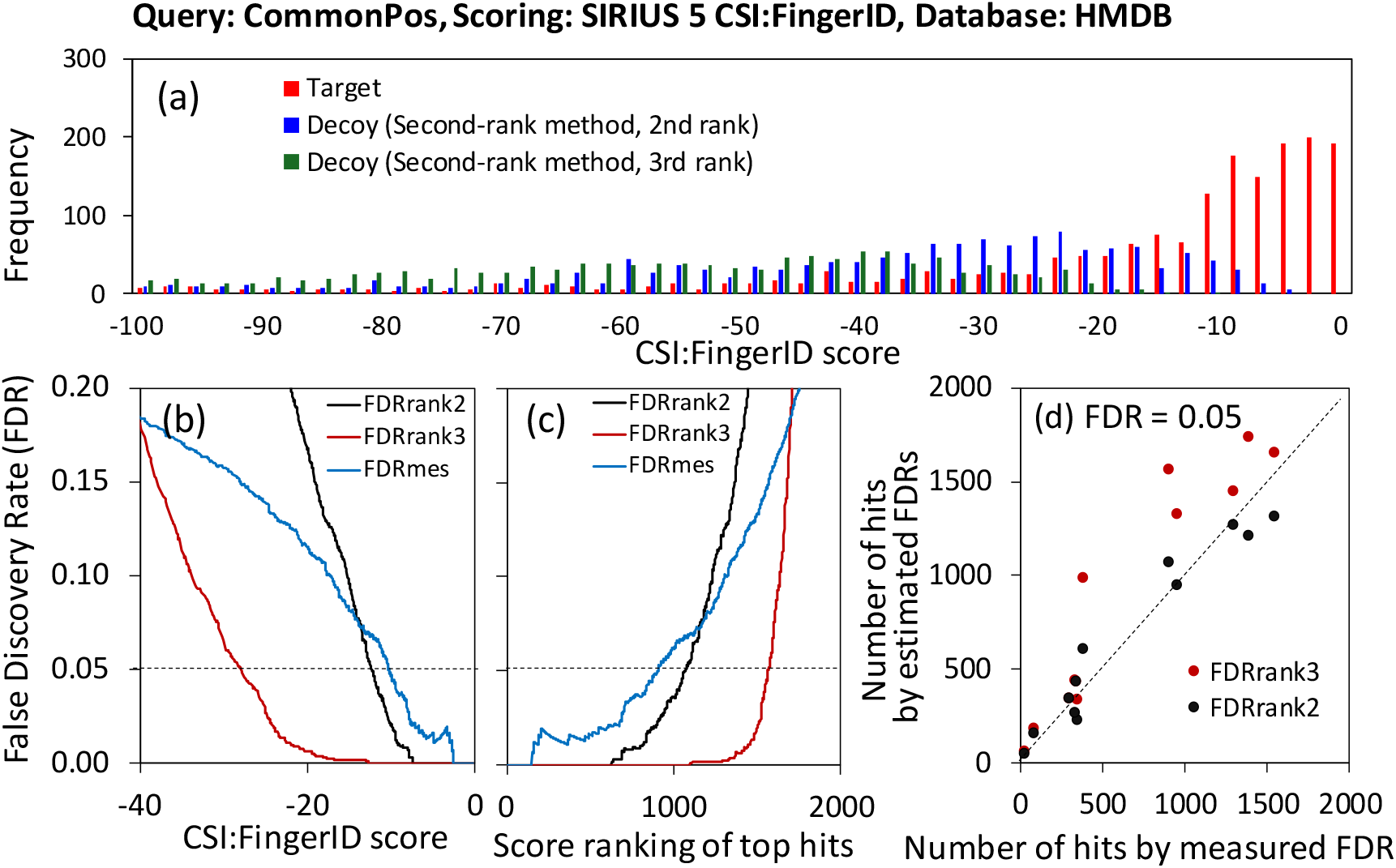
Performance evaluation of the second-rank method.(a) Distribution of the top-(Target), second- and third-ranked scores in the compound identification result. Total 3388 high-resolution mass spectra dataobtained at positive ion modewere collected from MassBank. The CommonPos dataset were served for the compound search by the SIRIUS 5 CSI:Finger ID scoring method using HMDB as the compound database. (b) Relationship between the CSI:Finger ID score and FDR levels. FDRmes indicate true FDR levels measured from the numbers of true and false hits (Fig. 2a). FDRrand2 and FDRs rank3 represent the estimated FDR (D/T) levels from the numbers of target (T) and decoy (D) hits in panel a. (c) Relationship between the score ranking of top hits and FDR levels. (e) Comparison between the number of hits determined by the true FDR (FDRmes) and estimated FDR (FDRrank2 and FDRrank3).

### Application to human metabolome DDA datasets

Previously, DDA metabolomic datasets were shown to contain similar product ion spectra obtained from the same compound^29)^. It has also been demonstrated that the signal-to-noise ratio of the product ion spectra can be improved by integrating similar spectra into one averaged spectrum^28)^. Four human metabolomics DDA datasets were obtained from a public repository. Four sets of averaged spectra were produced from the DDA datasets and used for the compound search by the SIRIUS 5 CSI:Finger ID scoring method, using the Biodatabase as the compound database. Furthermore, the pseudo-target-decoy and the 2nd rank method were used to control FDR = 0.05. The results are presented in Table 2. For Dataset 1, 3,242,674 product ion spectra across 248 data files were consolidated into 21,403 averaged spectra via spectral averaging. Among them, 8,104 spectra included three or more product ions and were used for the compound search. The number of hits at FDR=0.05, was estimated to be 709 and 642 by FDRc and FDRrand2, respectively, indicating that the second-rank method (FDRrand2) yielded more conservative results than the pseudo target-decoy method. The results for the other three datasets also showed that more conservative results were obtained using the second-rank method (FDRrand2).

**Table 2.**
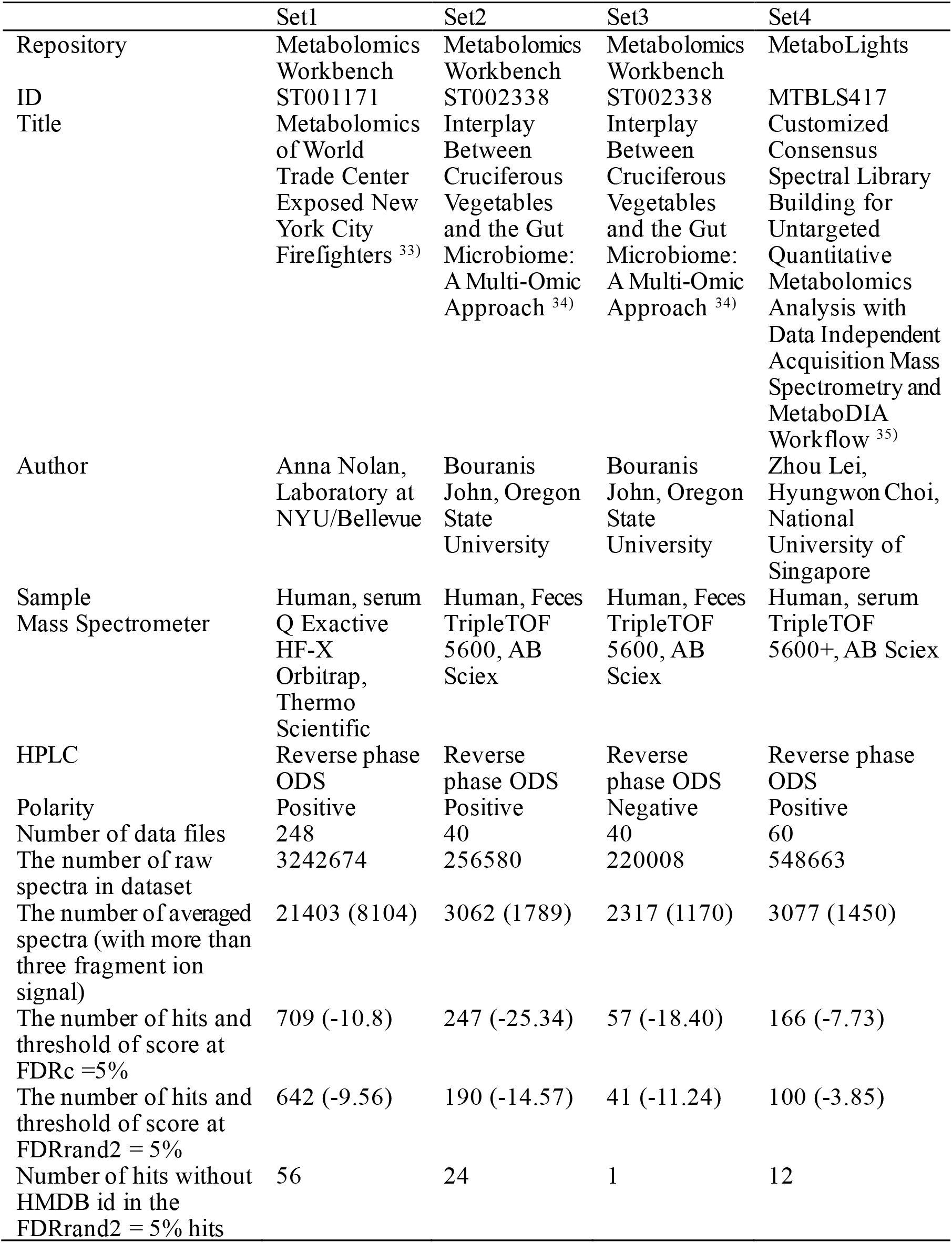
FDR based compound searching of human metabolome DDA datasets.

Next, the results of the compound search were investigated to identify metabolites not included in the human metabolome database (HMDB). Among the 642 hits of Dataset 1 estimated by FDRrand2, 56 hits did not have HMDB identifiers (Data S4). For example, the top hit for the spectrum shown in Figure 4a was *N*-lauroylethanolamide, whose score (−0.380) was highest among the 56 non-HMDB hits. The compound search result was confirmed using the measured spectrum of *N*-lauroylethanolamide in MassBank (MSBNK-BGC_Munich-RP003002; similarity score = 0.7437; data not shown). *N*-Lauroylethanolamide is a lipid mediator in mammals, indicating that its detection in human serum samples is plausible^30,31)^. Furthermore, for Dataset 2, the spectrum shown in Figure 4b hit to *β*-casomorphin 4 with the highest score (−1.01) among the 24 non-HMDB hits (Data S5). Although MassBank does not contain any measured spectra of this compound, the product ion spectrum was consistent with the possible fragmentation of *β*-casomorphin 4 by manual curation. *β*-casomorphin 4 is a tetprapeptide (Tyrosyl-prolyl-phenylalanyl-proline, Fig. 4c) and a degradation product of the milk protein *β*-casein, likely to be detected in human fecal samples^32)^.

**Figure 4.**
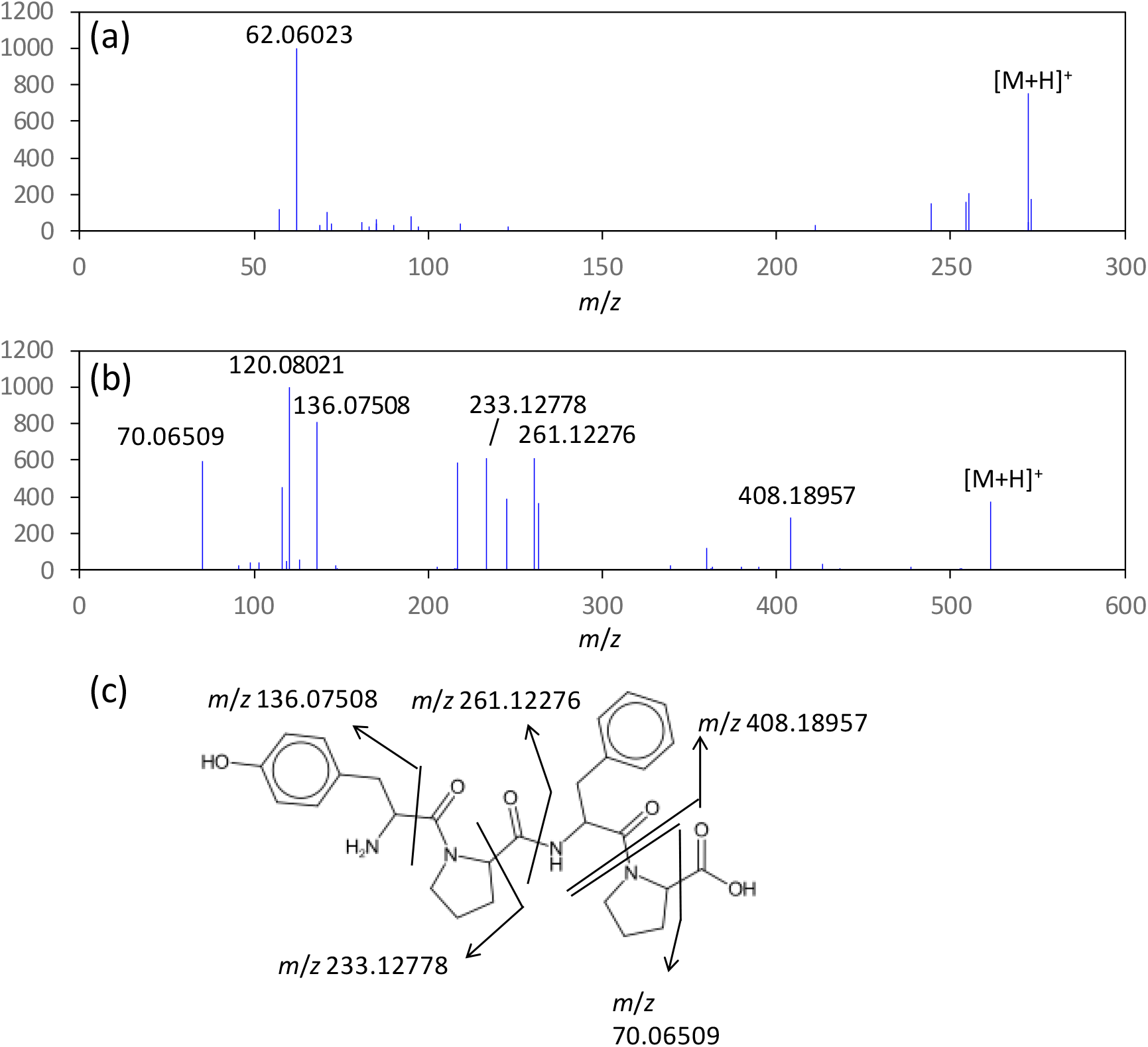
Two annotation examples with compounds not included in the human metabolome database (HMDB). (a) Product ion spectrum derived from dataset 1. The spectrum was annotated as *N*-lauroylethanolamide by the SIRIUS 5 CSI:Finger ID scoring method using Biodatabase as the compound database. (b) Product ion spectrum derived from dataset 2, which was annotated as *β*-casomorphin 4. (c) Structureof *β*-casomorphin 4 and estimated assignment of fragment ions.

## Discussion

In this study, the pseudo-target-decoy and second-rank methods were examined for the purpose of controlling the FDR in the compound search results of non-targeted metabolomics (Fig. 1). Notably, the calculated FDRs were approximate estimates because both methods were based on invalid heuristics or assumptions. In particular, estimations at FDR = 0.01 or 0.10, would contain extremely large errors (Fig 2 and 3). Even when estimating at an FDR of 0.05, it is necessary to adopt more conservative results to avoid underestimation. The results of this study revealed that the second-rank method (FDRrand2) provided more conservative results than the pseudo-target-decoy method (FDRc) (Fig. 2e and 3d). Using the two developed methods, a compound search was performed on four human metabolomic DDA datasets with FDR = 0.05 (Table 2). From the FDR controlled compound search results, we successfully found compounds not present in HMDB, such as *N*-lauroylethanolamide and *β*-casomorphin 4 (Fig. 4).

The two methods examined in this study offer the advantages of being simple and not requiring any random sampling techniques and modifications to structure elucidation software. However, they were approximate FDR estimation methods. In order to improve the estimation, the FDR estimation methods development for the spectra similarity search, such as spectra and fragmentation tree-based construction of decoy spectra^17)^, are promising for the construction of decoy query spectra datasets.

Nevertheless, the versatility of these methods is yet to be investigated. The software packages and their scoring methods are being developed at very rapid pace. Recently, SIRIUS 6 with an updated scoring method was opened to the public from the developer’s webpage (https://bio.informatik.uni-jena.de/software/sirius/). Furthermore, the product ion spectral data stored in MassBank were used for method verification because of their large structural variety. Additional verification is required to handle product ion spectra with less structural variation, such as in lipidomics. In the future, it is expected that more accurate FDR estimation can be achieved by developing methods based on more valid assumptions or methods for constructing valid decoy compound databases.

## Acknowledgment

We thank Profs. Yoshihiro Izumi at Kyushu University, Akiyoshi Hirayama at Keio University, Hiroshi Tsugawa at Tokyo University of Agriculture and Technology, Shujiro Okuda at Niigata University, and all Shin-MassBank project members for their helpful comments and support. This study was supported by the JST-NBDC Life Science Database Integration Project (grant number JPMJND2305).

---

**Figure S1.**
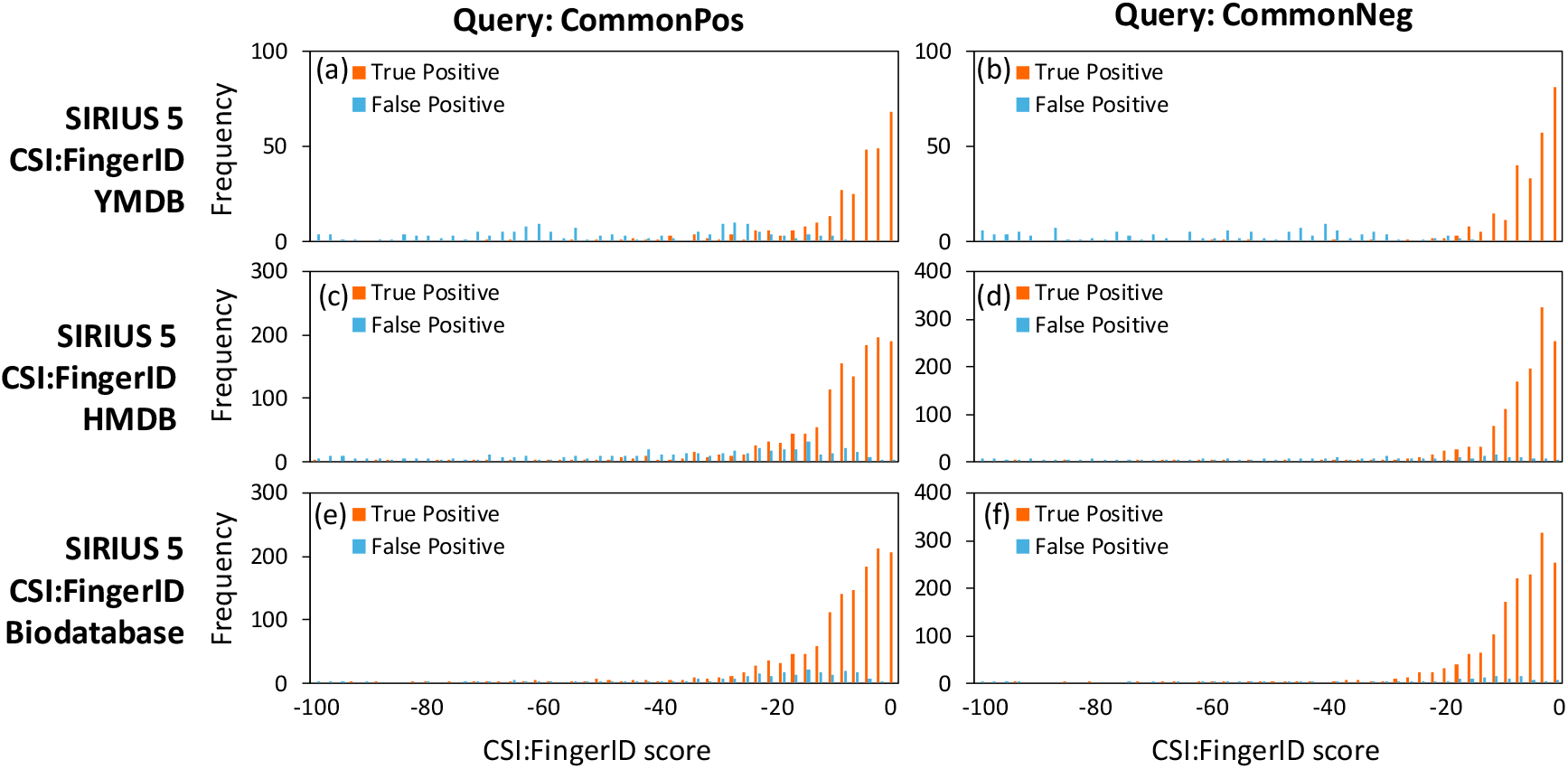
Score distribution of true positive and false positive hits in the compound identification result. Total 3388 (CommonPos) and 3100 (CommonNeg) high-resolution mass spectra data obtained at positive and negative ion mode were collected from MassBank, respectively. These datasets were served for the compound search by the SIRIUS 5 CSI:Finger ID scoring method using the three compound databases including (a,b) YMDB (c,d) HMDB, and (e,f) Biodatabase. Biodatabase is an integrated compound database available in SIRIUS 5.

**Figure S2.**
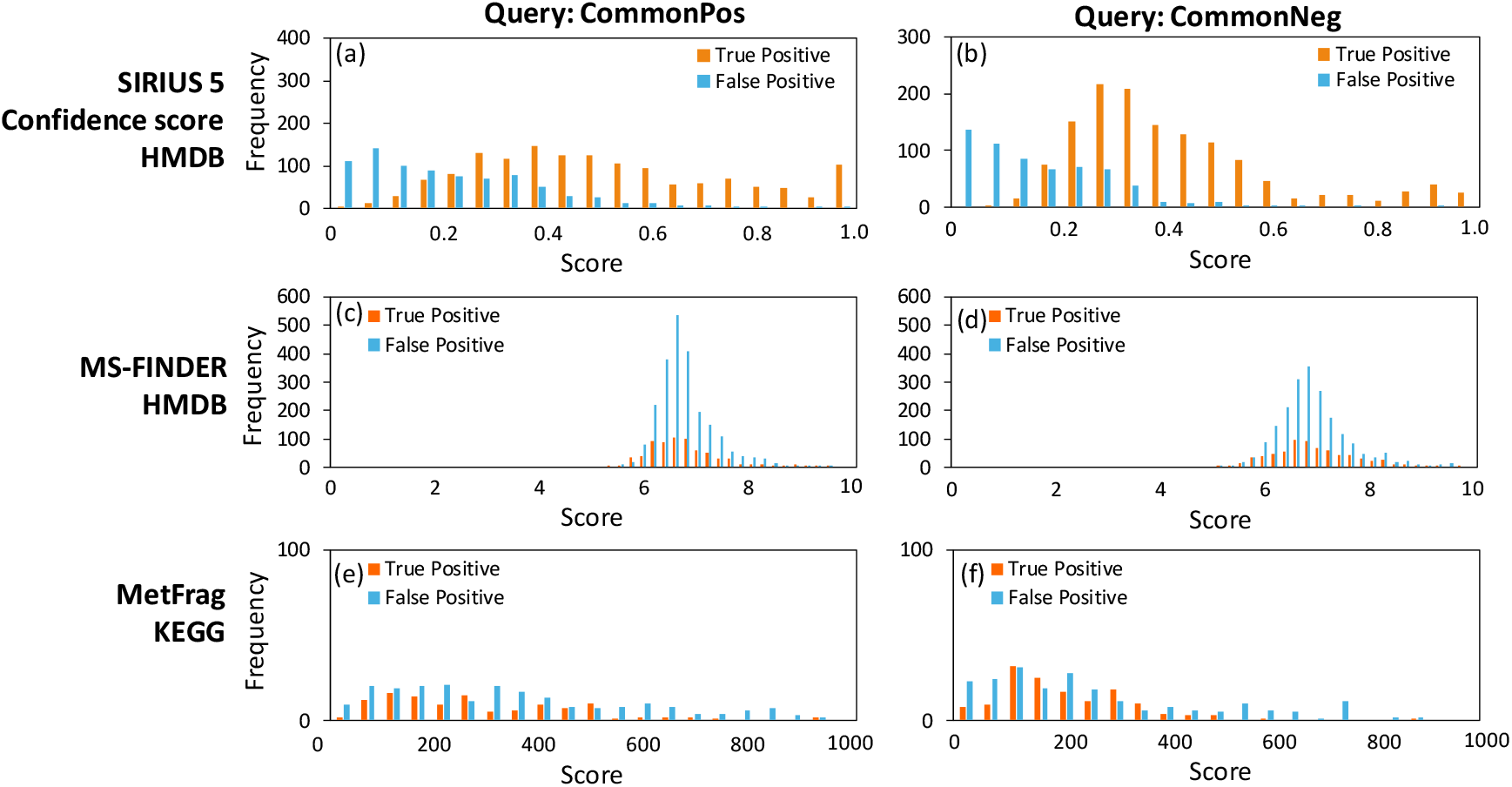
Score distribution of true positive and false positive hits in the compound identification result. Total 3388 (CommonPos) and 3100 (CommonNeg) high-resolution mass spectra data obtained at positive and negative ion mode were collected from MassBank, respectively. These datasets were served for the compound search by (a, b) SIRIUS 5 Confidence score, (c, d) MS-FINDER (c, d), and (e, f) MetFrag (e,f) using the HMDB and KEGG compound databases.

**Figure S3.**
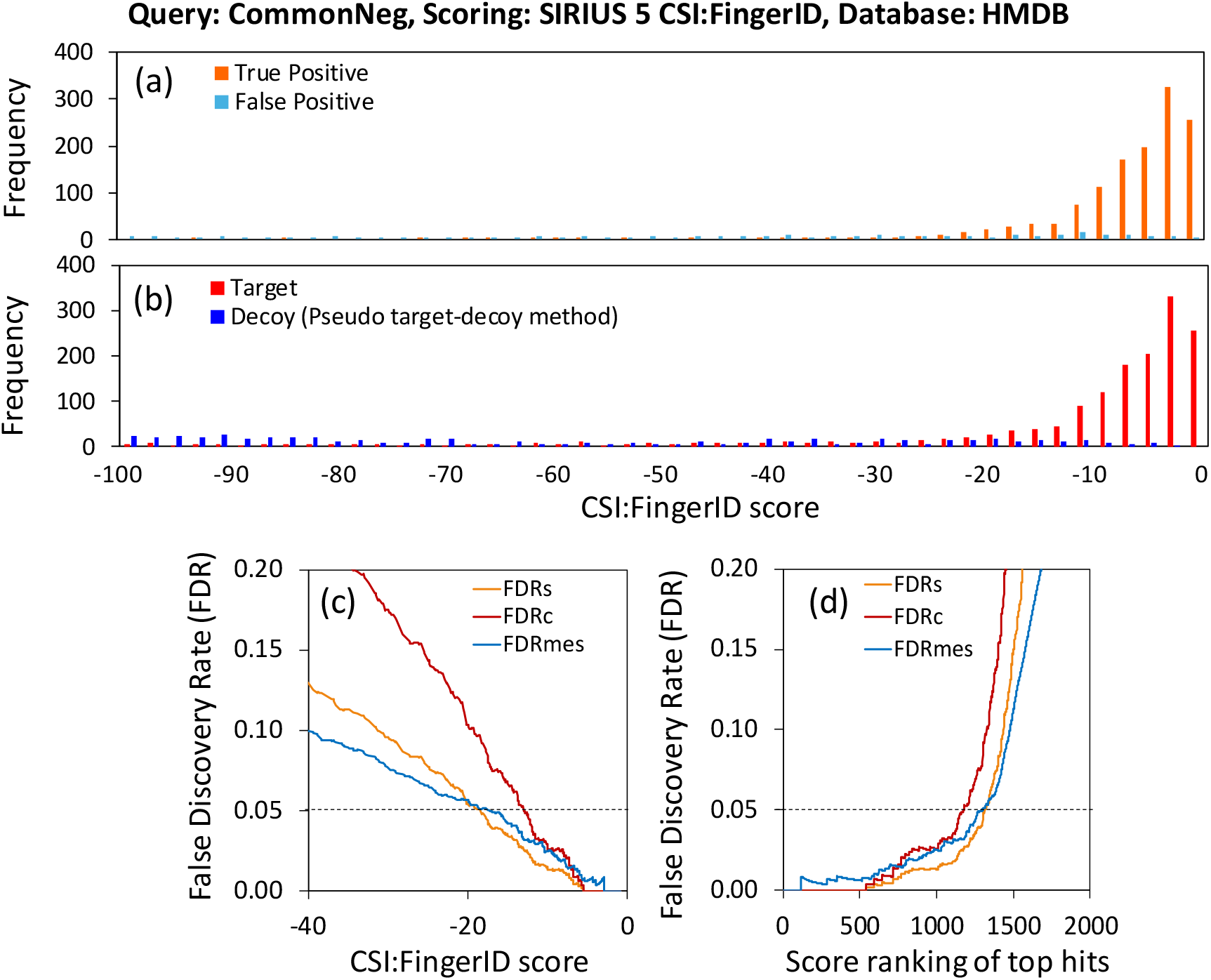
Performance evaluation of the pseudo-target-decoy method. (a) Score distribution of true positive and false positive hits in the compound identification result using the CommonNeg datasets. Total 3100 high-resolution mass spectra data obtained at negative ion mode were collected from MassBank. The CommonNeg dataset were served for the compound search by the SIRIUS 5 CSI:Finger ID scoring method using HMDB as the compound database. (b) Score distribution of searching results of target and decoy of CommonNeg datasets. (c) Relationship between the CSI:Finger ID score and FDR levels. FDRmes indicate true FDR levels measured from the numbers of true and false hits in panel a. FDRs and FDRs represent estimated FDR levels from the numbers of target (*T*) and decoy (*D*) hits in panel b. FDRs = *D*/*T*, FDRc = 2*D*/(*T*+*D*) (d) Relationship between the score ranking of top hits and FDR levels.

**Figure S4.**
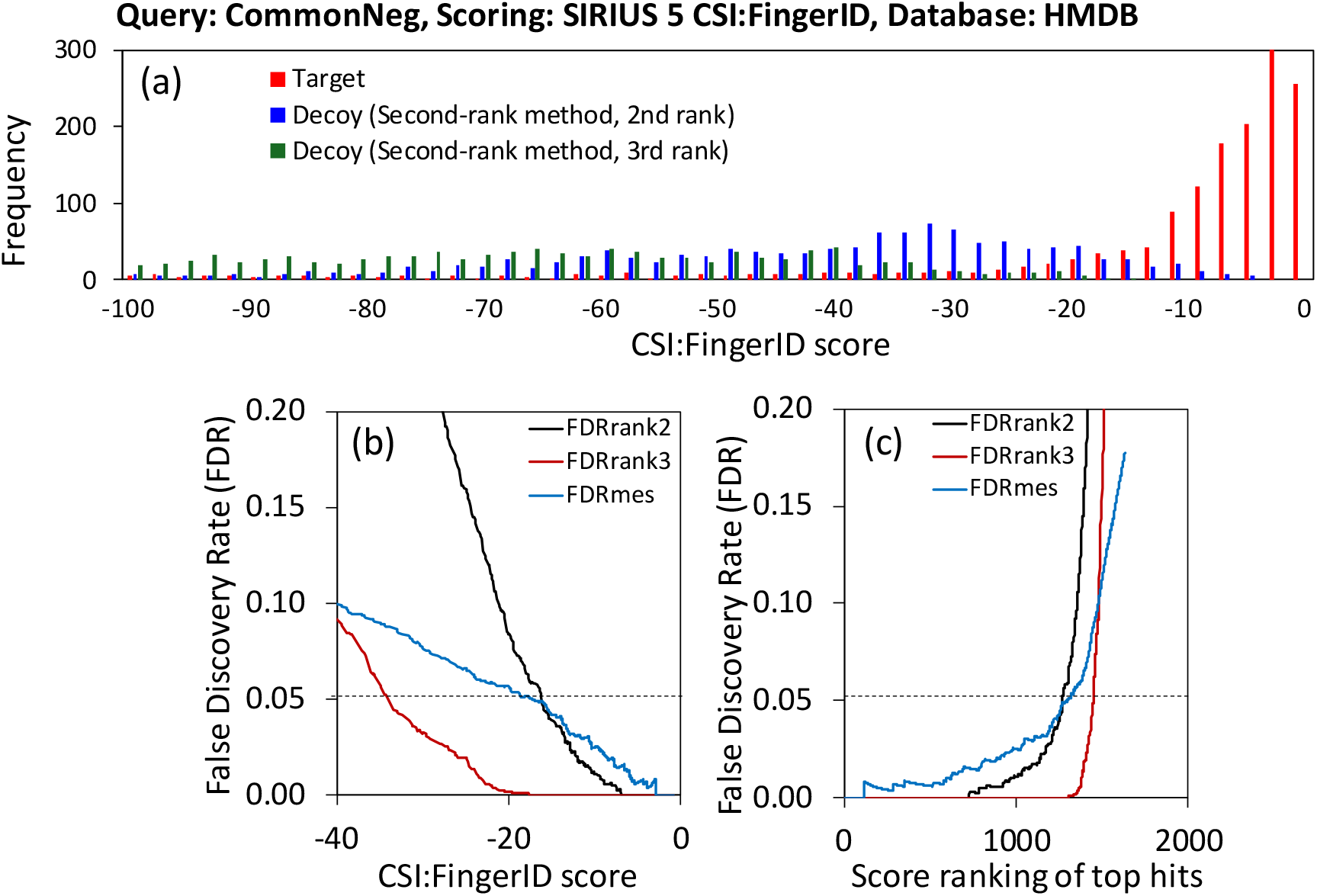
Performanceevaluation of the second-rank method.(a) Distribution of the top-(Target), second- and third-ranked scores in the compound identification result. Total 3100 high-resolution mass spectra data obtained at positive ion mode were collected from MassBank. The CommonNeg dataset were served for the compound search by the SIRIUS 5 CSI:Finger ID scoring method using HMDB as the compound database.(b) Relationship between the CSI:Finger ID score and FDR levels. FDRmes indicate true FDR levels measured from the numbers of true and false hits (Fig. S3a). FDRrand2 and FDRs rank3 represent estimated FDR (*D*/*T*) levels from the numbers of target (*T*) and decoy (*D*) hits in panel a. (c) Relationship between the score ranking of top hits and FDR levels. (e) Comparison between the numbers of hits determined by the true FDR (FDRmes) and by the estimated FDR (FDRrank2 and FDRrank3).

**Table S1a.**
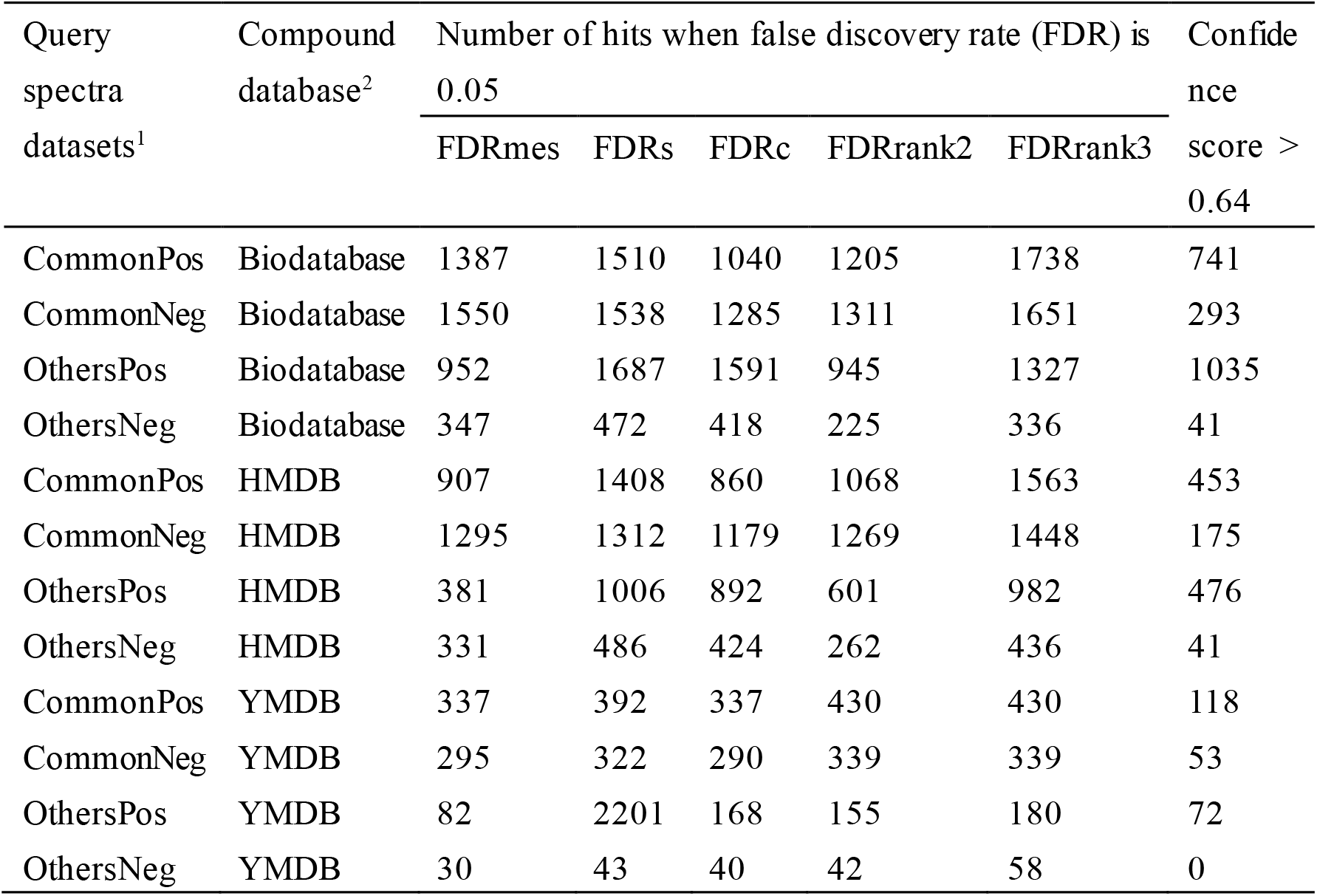
(a) Number of hits in the compound searching results by SIRIUS 5 CSI:Finger ID scoring method when false discovery rate (FDR) is 0.05

**Table S1b.**
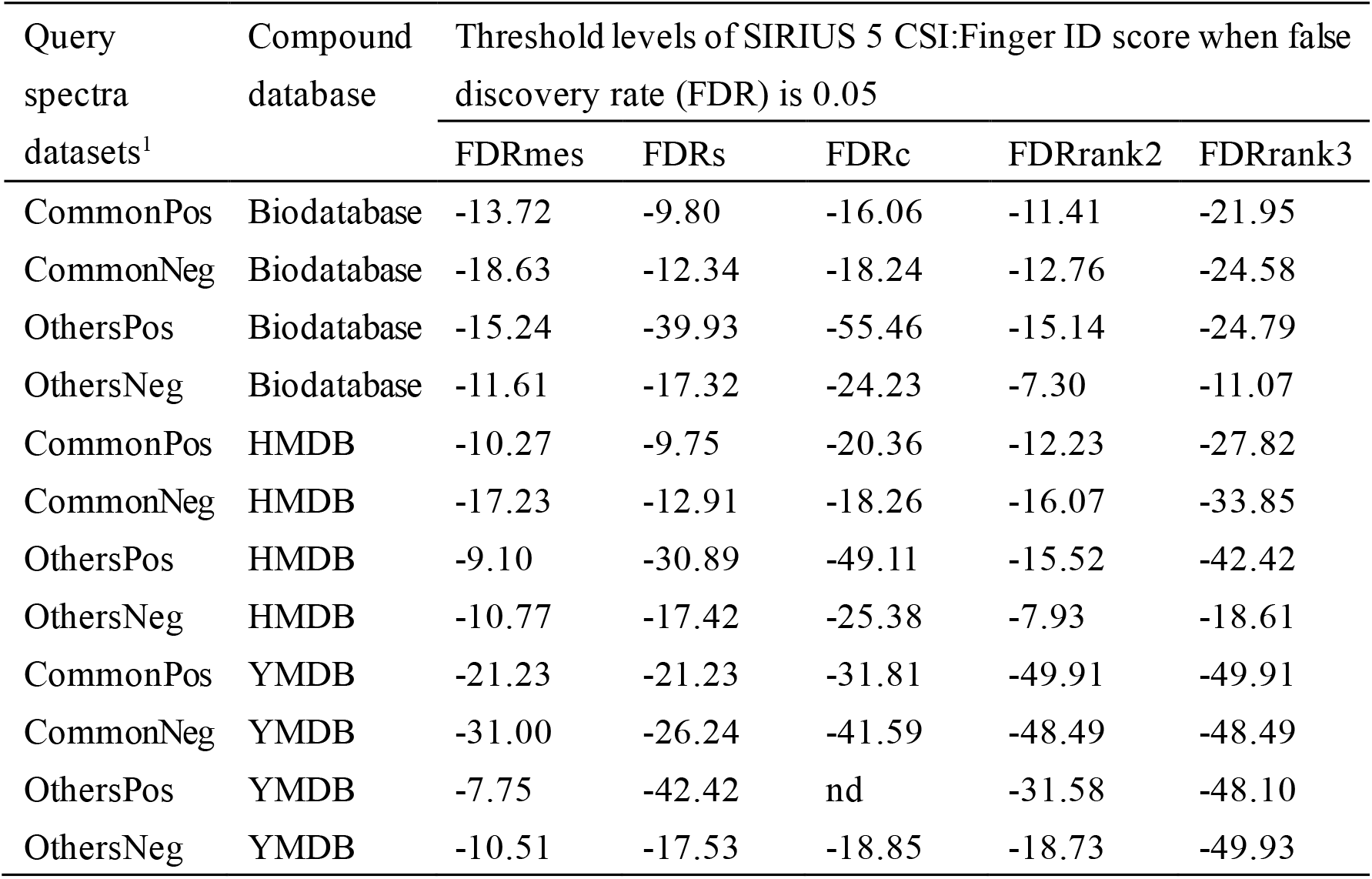
(b) Threshold levels of SIRIUS 5 CSI:Finger ID score when false discovery rate (FDR) is 0.05

## Notes

### Competing Interest Statement

The authors have declared no competing interest.

